## Supplemental Figures for "Genome-wide off-rates reveal how DNA binding dynamics shape transcription factor function"

### Supplemental Figure 1

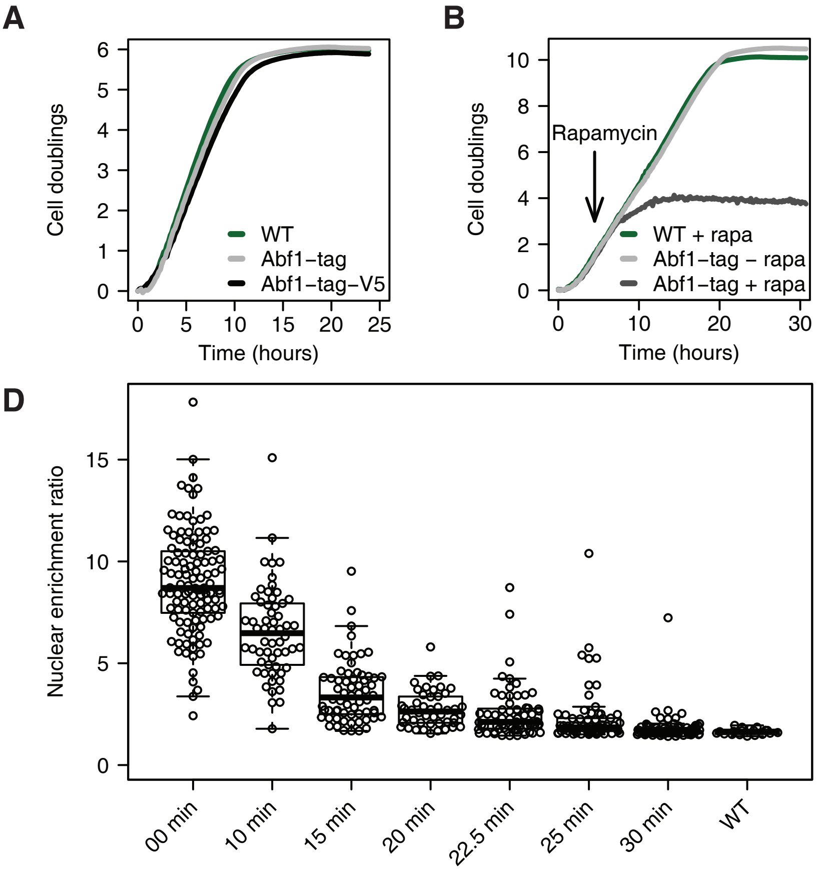


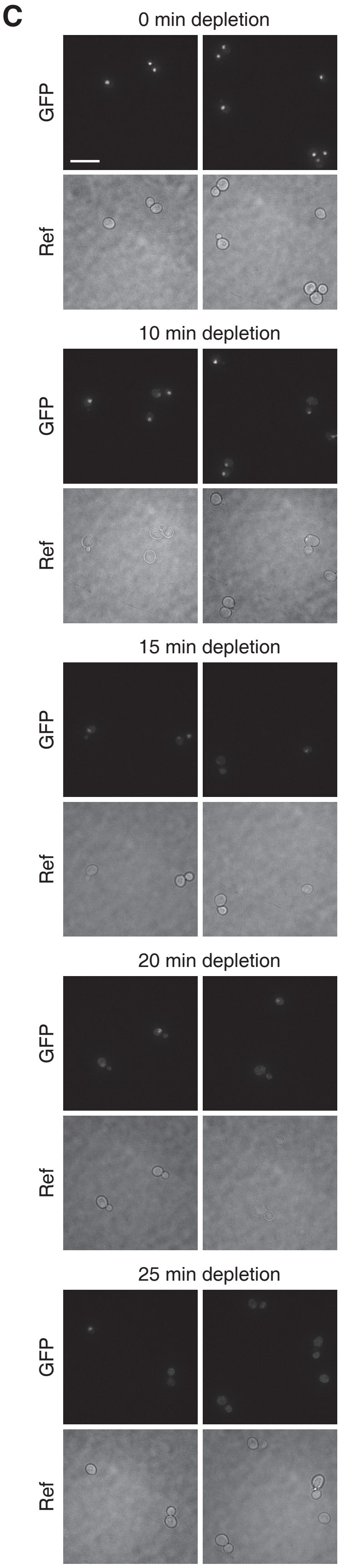


**Supplemental Figure S1. Dynamics of Abf1 depletion**
**(A)** Representative growth curves of the untagged parental background strain (WT), an ABF1 FRB-GFP-tagged strain (Abf1-tag) and an ABF1 FRB-GFP-V5-tagged strain (Abf1-tag-V5). The WT strain is the parental BY4742/S288C strain of the Abf1 anchor-away strains that have been genetically desensitized to rapamycin. The Abf1-tag strain grows approximately 10% slower compared to the WT strain and the Abf1-tag-V5 strain grows approximately 15% slower than the WT strain. **(B)** Growth curves of the WT strain with rapamycin, and the ABF1 FRB-GFP tagged strain with and without rapamycin, the inducing agent for nuclear depletion (Haruki *et al*, 2008). Rapamycin was added 4.5 hours after the start of the experiment to induce nuclear depletion (arrow). Upon induction of depletion, clear disruption of growth is visible after approximately 3 hours ($t$ = 7.5h) and cessation of growth is visible after approximately 8 hours ($t$ = 12.5h). **(C)** Representative fluorescence microscopy images of ABF1 FRB-GFP-V5 cells before (0 minutes) and at several time points after (10, 15, 20 and 25 minutes) induction of nuclear depletion. Scale bar: 10 μm. **(D)** Quantification of nuclear enrichment ratios of ABF1 FRB-GFP and ABF1 FRB-GFP-V5 cells during depletion (three biological replicates). Both DNA-bound and free Abf1 contribute to the GFP signal and therefore the decrease in fluorescence represents the depletion speed of both unbound as well as bound Abf1. Dots represent individual cell measurements. The nuclear enrichment ratio was calculated as the maximum intensity divided by the median intensity of each cell. A WT strain without GFP was taken along as a control.

### Supplemental Figure 2

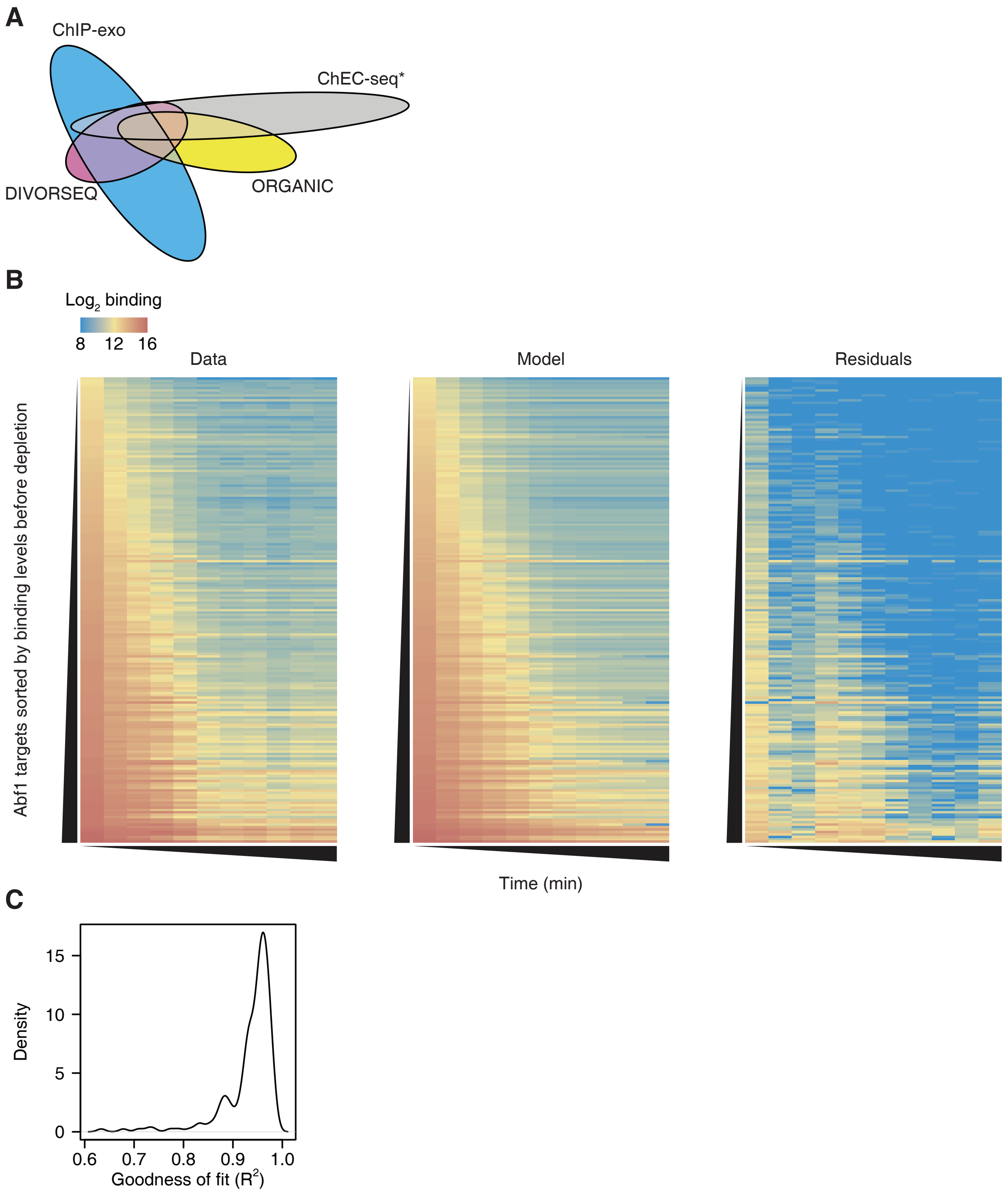


**Supplemental Figure S2. The exponential decay model closely fits the binding data**
**(A)** Venn diagram showing the overlap between all unfiltered Abf1 binding sites detected here (*n*=948) and published Abf1 binding sites detected using ORGANIC ChIP (Kasinathan et al, 2014; *n*=1068), binding sites detected using ChEC-seq that were annotated both as "fast" and "high scoring" (Zentner et al, 2015; *n*=1583) and binding sites that were detected using ChIP-exo v5 (Rossi et al, 2018b; *n*=3177). **(B)** Heatmap representation of the average Abf1 binding at the 191 binding sites, quantified as the $\log_2$ number of reads in the peak (peak summit ± 50 bp). The rows are sorted on the binding levels before depletion, and the columns represent increasing time of depletion. From left to right: average binding data, fit and residuals of the fit (median absolute deviation) are shown. **(C)** The distribution of the goodness of fit of the 191 Abf1 binding sites.

### Supplemental Figure 3

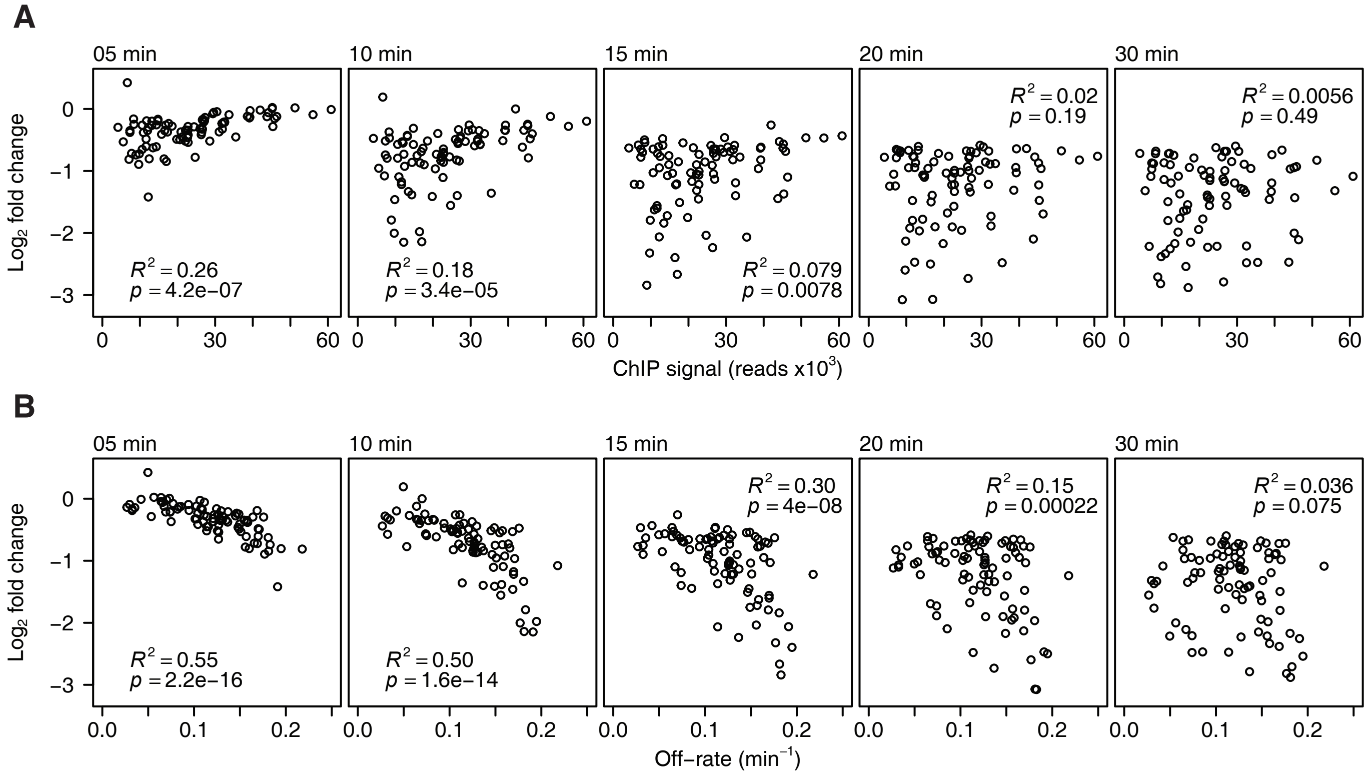


**Supplemental Figure S3. Changes in mRNA synthesis correspond more closely to off-rates than binding levels** **(A)** Change in mRNA synthesis rate versus Abf1 binding level before depletion of the corresponding binding site after 5, 10, 15, 20 and 30 minutes of depletion of downregulated genes (fold change > 1.5 and $p < 0.01$ at 20 and 30 minutes of depletion) with Abf1 binding in the promoter ($n$=88). The panel for $t$ = 10 min is also shown in **Fig 3E**. **(B)** Change in mRNA synthesis rate versus the off-rates of the corresponding binding sites for the same gene-binding sites pairs as in (A) 5, 10 ,15, 20 and 30 minutes after induction of depletion. The panel for $t$ = 10 min is also shown in **Fig 3F**.
